# CETP inhibitor evacetrapib enters mouse brain tissue

**DOI:** 10.1101/2023.02.21.529381

**Authors:** Jasmine Phénix, Jonathan Côté, Denis Dieme, Sherilyn J Recinto, Felix Oestereich, Sasen Efrem, Sami Haddad, Michèle Bouchard, Lisa Marie Munter

**Affiliations:** Department of Pharmacology and Therapeutics, McGill University, 3649 Promenade Sir William Osler, Montreal, Quebec, Canada H3G 0B1; Department of Environmental and Occupational Health, School of Public Health, Université de Montreal, Montreal, Quebec, Canada; Public Health Research Center (CReSP), Université de Montréal, Montreal, Quebec, Canada; Cell Information Systems group, Bellini Life Sciences Complex, 3649 Promenade Sir William Osler, Montreal, QC, Canada H3G 0B1; Integrated Program in Neuroscience, McGill University, Montréal, QC, Canada H3A 2B4; Centre de Recherche en Biologie Structurale (CRBS), 3649 Promenade Sir William Osler, Montreal, QC, Canada H3G 0B1

## Abstract

High levels of plasma cholesterol, especially high levels of low-density lipoprotein-cholesterol (LDL-C), have been associated with an increased risk of Alzheimer’s disease. The cholesteryl ester transfer protein (CETP) in plasma distributes cholesteryl esters between lipoproteins and increases LDL-C in plasma. Epidemiologically, decreased CETP activity has been associated with sustained cognitive performance during aging, longevity, and a lower risk of Alzheimer’s disease. Thus, pharmacological CETP inhibitors could potentially be repurposed for the treatment of Alzheimer’s disease as they are safe and effective at lowering CETP activity and LDL-C. While CETP is mostly expressed by the liver and secreted into the bloodstream, CETP is also expressed by astrocytes in the brain. It is therefore important to determine if CETP inhibitors can enter the brain. Here, we describe pharmacokinetic parameters of the CETP inhibitor evacetrapib in plasma, liver, and brain tissues in CETP transgenic mice. We show that evacetrapib crosses the blood-brain barrier and is detectable in brain tissue 0.5 h after a 40 mg/kg i.v. injection in a nonlinear function. We conclude that evacetrapib may prove to be a good candidate to treat CETP-mediated cholesterol dysregulation in Alzheimer’s disease.

## Introduction

Cholesterol plays major roles in diverse processes essential to life and is ubiquitously found in membranes and organelles in mammalian cells (Maxfield and van Meer, 2010; Paschkowsky et al., 2018). Cholesterol acts as a signal transducer for many signaling pathways and is the precursor for steroid hormones (Burke et al., 1999; Cooper et al., 2003; Payne and Hales, 2004; Porter et al., 1996). Though cholesterol is necessary for proper development and maintaining homeostasis, dysregulated cholesterol metabolism has also been shown to be associated with a plethora of pathologies from cardiovascular diseases to neurodegenerative diseases including Parkinson’s disease, amyotrophic lateral sclerosis, and Alzheimer’s disease (Godoy-Corchuelo et al., 2022; Huang et al., 2019; Iwagami et al., 2021; Whitmer et al., 2005). Interestingly, higher levels of plasma cholesterol and LDL-C were also associated with an increased risk of developing Alzheimer’s disease (Liu et al., 2020; Solomon et al., 2009; Whitmer et al., 2005). However, the molecular mechanisms by which peripheral LDL-C may increase the risk for Alzheimer’s disease remain unclear, especially since LDL particles do not cross the blood-brain-barrier (Bjorkhem and Meaney, 2004).

The cholesteryl ester transfer protein (CETP) is a plasma glycoprotein mostly secreted by the liver, spleen, adipose tissues, and brain (Tall, 1995). Its main function is to mediate the transport of cholesteryl esters and triglycerides between plasma lipoproteins. The CETP transporter works according to a concentration gradient leading to a net flux of cholesteryl esters from high-density lipoproteins (HDL) particles into LDL and very low-density lipoproteins (VLDL) increasing the total LDL-C and VLDL-C levels (Barter and Rye, 1994). Thus, CETP activity is proposed to increase the risk of AD through raising LDL-C. CETP inhibitors were proposed as a new class of non-statin cholesterol-lowering drugs to increase HDL-C and decrease LDL-C in plasma such as dalcetrapib (Hoffman-La Roche) (Ray et al., 2014), anacetrapib (Merck) (Gotto et al., 2014), evacetrapib (Eli Lilly) (Nicholls et al., 2017), or obicetrapib (Dezima Pharma) (Nicholls et al., 2022; Tall and Rader, 2018). These, new generation CETP inhibitors were demonstrated to be safe and proven to be effective at lowering LDL-C while raising HDL-C to varying extents. However, they failed to reduce cardiovascular events over already existing drugs and thus were not approved for the treatment of cardiovascular disease (Nelson et al., 2022; Tall and Rader, 2018). Nevertheless, CETP inhibitors may carry potential for being repurposed in other conditions such as Alzheimer’s disease.

CETP forms a promising drug target since individuals with homozygous CETP deficiency have been described as overall healthy with potentially longer life span (Brown et al., 1989; Inazu et al., 1990). Remarkably, the CETP polymorphism rs5882 (I405V) was associated with exceptional longevity and healthy aging in centenarians (Barzilai et al., 2006; Barzilai et al., 2003). This CETP variant and others that were subsequently identified lead to either decreased CETP expression or decreased CETP activity, and associate with good cognitive performance and decreased risk of developing Alzheimer’s disease (Barzilai et al., 2003; Lythgoe et al., 2015; Sun et al., 2013). Here, it is important to note the role of the apolipoprotein E isoform 4 (APOE4), a protein important for the cellular uptake of LDL particles. APOE4 is the greatest genetic risk factor for Alzheimer’s disease increasing the risk by 3-15 fold in a dose-dependent manner (Bellenguez et al., 2022; Neu et al., 2017; Poirier et al., 1993). CETP variants with lower CETP activity have the potential to decrease Alzheimer’s disease risk specifically in subjects carrying the APOE4 allele (Chen et al., 2008; Oliveira and de Faria, 2011; Rodriguez et al., 2006; Sundermann et al., 2016). Thus, pharmacological CETP inhibition may have the capacity to abolish the increased Alzheimer’s risk conferred by APOE4, again indicating that CETP is a valuable drug target with promising disease-modifying outcomes. As such, the use of CETP inhibitors may positively impact cognitive performance, promote longevity, and decrease the risk of Alzheimer’s disease.

To evaluate the potential of CETP inhibitors for Alzheimer’s disease prevention, preclinical modeling in mice is necessary. Of note, mice do not express CETP or any orthologous genes and as a result, mice have naturally very low LDL levels, thereby limiting the use of such rodents in investigating LDL-related diseases. Therefore, transgenic mice expressing the human *CETP* gene (*hCETPtg*) were developed in 1991 and have since widely been used in cardiovascular disease research (Agellon et al., 1991; Jiang et al., 1992). hCETPtg mice have demonstrated a similar CETP expression pattern to humans, and most importantly, on a high cholesterol diet they show elevated LDL plasma levels comparable to those of healthy individuals(Steenbergen et al., 2010). In humans, CETP is mostly expressed in the liver and secreted to the plasma but has also been found in the cerebrospinal fluid at 12% of the concentration found in the plasma, as well as in the brain. As plasma CETP does not cross the blood-brain barrier, this suggests direct synthesis of CETP within the brain (Albers et al., 1992; Yamada et al., 1995). Indeed, our laboratory previously confirmed *CETP* mRNA expression in astrocytes enriched from hCETPtg mouse brains (Oestereich et al., 2022). The functions carried out by cerebral CETP remain unclear, since only HDL-like particles are formed in the brain and there are no LDL-like particles. Thus, the question as to whether CETP of the brain or of the periphery modulates the risk of Alzheimer’s disease prevails. To first assess if the potent CETP inhibitor evacetrapib can reach brain tissue, we herein characterized the pharmacokinetics of evacetrapib in hCETPtg mice.

## Materials and Methods

### Chemicals

Evacetrapib was purchased from Adooq Biosciences (USA). All other reagents were commercially available and were pure or liquid chromatography/mass spectrometry (LC/MS) grade.

### Animal acclimatation and housing

hCETPtg mice with a C57BL/6J background were chosen as the experimental model (Jackson Laboratory). Mice were housed in the Goodman Cancer animal facility (12-h light-dark cycle; 20-26°C and 40-60% humidity) in polysulfone cages of two to five mice per cage. Husbandry was heterozygous and mice were genotyped by Transnetyx^TM^, at 3 weeks of age. The protocol was approved by the Animal Compliance Office from the McGill University (ACO approval# 2013-7359).

### Animal exposure and sample collection

The experiment was carried out in conformity with the OECD Guidelines 417(OECD, 2010). Male hCETPtg mice were chosen for the pharmacokinetic analysis. Mice were fed Teklad Global Diets #2018 from Envigo, Canada, for 8 weeks starting from birth followed by a diet change of Modified Low-Fat RD Western diet to match TD.08485 with 1% cholesterol from Research Diets #D16121201 for a duration of three to four weeks. Throughout that period, tap water was provided *ad libitum*. Initially, evacetrapib was solubilized in 20% of the final solution volume of Kolliphor® EL (pH-range 6.0-8.0 cat# C5135) by alternating between 5-minute intervals using an ultrasonic water bath at 4°C and hot water bath at 50°C until complete dissolution. Then, 80% of the final volume was added (isotonic glucose solution 50 g/L glucose containing D-(+)-glucose Sigma cat# G7021 in MilliQ water, sterile filtered).

CETP inhibitor evacetrapib was administered through tail vein injection at doses of 40 mg/kg bodyweight (BW) or 120 mg/kg BW. Mice weighed 28.42 ± 3.16 g on the day of injections. Mice were randomly divided into two groups: 24 mice received a dose of 40 mg/kg BW of evacetrapib while 24 other mice received a dose of 120 mg/kg BW. Mice from both groups were sacrificed at 0, 0.5, 2, 6, 12, and 24 h post injection. Mice sacrificed at later time points had continued access to food and water until the end of the experiment. Blood was collected (300-500 µL) at sacrifice for each mouse by cardiac puncture. Blood plasma was extracted using 10 µL of 1 M EDTA pH 8.0 solution in MilliQ water. Brain and liver tissues were collected immediately after sacrifice, rinsed in 0.9% NaCl solution, and snap-frozen in liquid nitrogen. Samples were stored at -80°C until analysis.

### UHPLC-MS-qTOF analysis

Sample preparation and digestion for evacetrapib analysis was performed by UHPLC-MS-qTOF. Working solutions from a stock of evacetrapib were prepared at concentrations of 10 µmol, 100 nmol, 1 nmol, 500 pmol, and 125 pmol per mL of methanol. Working solutions of anacetrapib used as an internal standard was prepared at concentrations of 10 µmol, 100 nmol, 1 nmol per mL from a stock solution.

Brain and liver samples were homogenized using a polytron supplemented with a carbonate buffer at pH 9.8. The volume was adjusted such that each sample reached a final concentration of 100 mg/mL of buffer. A volume of 0.5 mL of homogenate was transferred to a pyrex tube and enriched with 100 µL of 1 nmol/mL anacetrapib solution used as an internal standard. Then, samples were extracted twice using 4 mL of saturated ethyl acetate. For each extraction, samples were shaken for 30 min using a lateral shaker followed by a centrifugation round (20 min, 3000 rpm, 4°C). Organic phases were collected and combined, then evaporated until dry under nitrogen stream in a rotating bath at 40°C. The residues were resuspended in methanol. Samples were vortexed until no residues were left. Resuspended residues for both brain and liver were then centrifuged (20 min, 3000 rpm, 4°C). The supernatant was recovered, transferred into new vials, and injected into the UHPLC-MS-qTOF. For the plasma measurement, 20 µL of plasma was enriched with 100 µL of 1 nmol/mL anacetrapib solution in a microtube. Evacetrapib was extracted using two steps of ethanol wash (200 µL) followed by an incubation time (30 min, 22°C) and centrifugation (5 min, 16000g, 10°C). To separate the liquid phase from the solid phase, samples were dried under a gentle nitrogen stream. The residues were subsequently dissolved in methanol, vortexed until no residues were left, and centrifuged (20 min, 3000 rpm, 4°C). The supernatant was recovered, transferred into new vials, and injected into the UHPLC-MS-qTOF.

The UHPLC-MS-qTOF system consisted of an Agilent model 1290-LC binary gradient UHPLC system (Agilent, Mississauga, Canada) connected to an Agilent model 1290 autosampler and thermostated column compartment (Agilent, Mississauga, Canada) and coupled to a quadrupole time-of-flight mass spectrometer with a Dual J et Stream Electrospray Ionization (Dual-AJS ESI) source (Agilent Technologies, Mississauga, Ontario, Canada). The AJS ESI interface was operated in positive ion mode. The column used was an Agilent Zorbax Eclipse plus C18 (2.1 × 50 mm; 1,8 µm). The precolumn used was Agilent *Fast Guard* Zorbax Eclipse plus C18 (2.1 X 5 mm; 1,8 µm). HPLC elution solutions were H_2_O eluant 0.01% acidified (1 L ultra-pure water + 100 µL of HPLC-grade acetic acid) and methanol eluant 0.01% acidified (1 L MS-grade methanol + 100 µL HPLC-grade acetic acid). Tuning of the instrument was performed once a month and a calibration was performed once a week to accurately analyze the mass ratio m/z < 1700.

The exact mass of evacetrapib was determined in MS mode using the following ToF conditions: sheath gas (N2) temperature at 365°C and gas flow rate of 10 L/min; nebulizer gas pressure of 50 psi; drying gas temperature (N2) at 200°C and flow rate of 12 L/min; capillary voltage (Vcap) at 3000 V, nozzle voltage at 1000 V, fragmentor at 75 V, skimmer at 65 V and octopole at 750 V. The precursor ion [M+H]^+^ evacetrapib of analyzed was m/z 639.28797. Identification and quantification were performed in Extracted Ion Chromatogram (EIC) mode.

Quantification was carried out using a seven-point-calibration curve with internal standard correction, prepared respectively in brain, liver or in plasma. These curves were established by plotting the response factors as a function of the concentration levels, over a maximum range of 6.25 to 200 pmol/mL. The response factors corresponded to the peak-area ratios of evacetrapib ion to the anacetrapib internal standard ion. The limit of detection (LOD) (corresponding to 3 times of standard deviations of response ratio from replicate analysis of a blank) were 30.3 pmol/g of brain, 11.5 pmol/g of liver and 83 pmol/mL of plasma. Repeatability from replicate analysis of samples under the same calibration and tuning conditions (blank samples spiked with evacetrapib standards at two levels) was below 5%. The recovery percentages determined in the different matrices with two levels of spikes were between 95 and 108%.

### Data analysis and determination of main kinetic parameters from different organ time courses

The time courses of evacetrapib in blood, brain, and liver were established following injection of 40 and 120 mg/kg BW in mice (expressed as the average concentration in nmol/mL or nmol/g over time). The parameters used to determine the fit come the following equation A_1_e^(b^_1*t)_ + A_2_e^(b^_2*t)_ where A_1_ and A_2_ are, the different intercepts within the function and b_1_ and b_2_ are the different slopes within the function, and t is the specific time for which the parameter is calculated. The concentration-time course data for each tissue was independently fitted with Microsoft Excel using the solver function set on the generalized reduced gradient (GRG) nonlinear method. Using the parameters generated by the fit, the slope and intercepts were determined. The elimination rate constant k_i_ and the half-life (t_1/2_) for each tissue at both concentrations were calculated using the following equations:

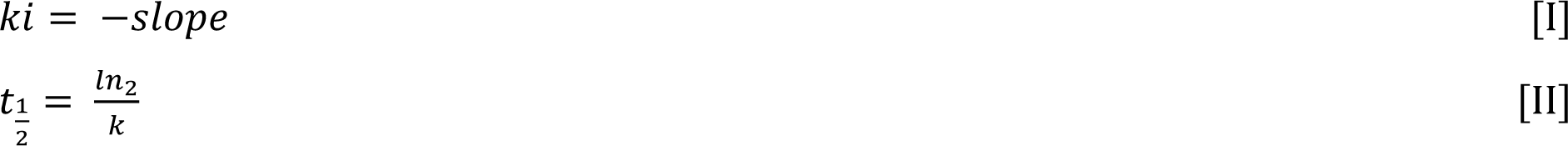

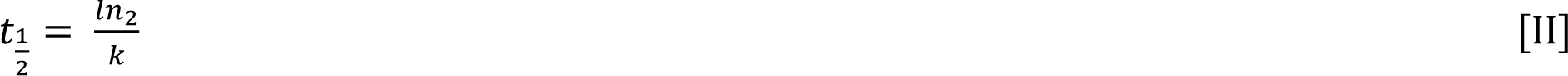

Pharmacokinetic parameters such as area under the plasma concentration-time curve (AUC_IV_), area under the first moment of the plasma concentration-time curve from time zero to infinity (AUMC_IV_), mean residence time (MRT), apparent total body clearance of the drug from plasma (CL_blood_) and apparent volume of distribution at steady state (Vss) were determined using the following equations:

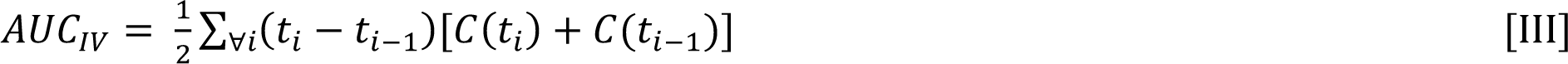

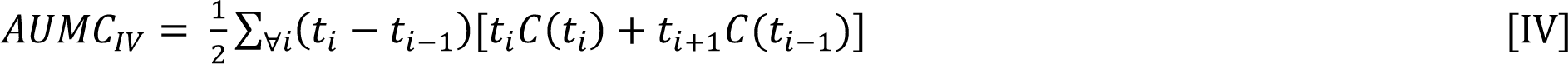

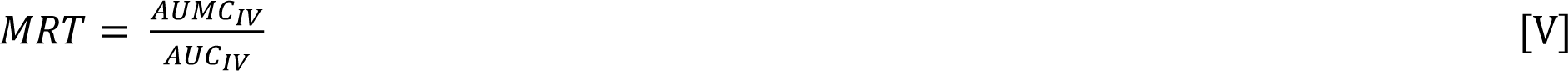

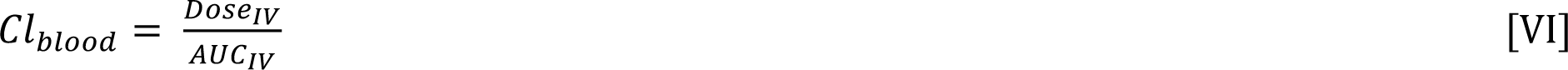

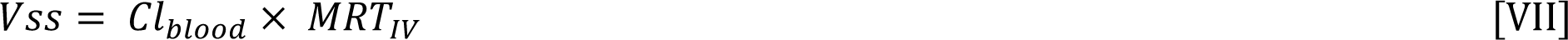

Mice were not perfused at sacrifice to allow for plasma, brain, and liver to be collected accurately at time point 0 and uniformly across all time points. Thus, residual blood in vessels of the brain may have left residual traces of evacetrapib in brain samples. Residual plasma volume in mouse brains was determined to be on average 8 µL according to a study by Kaliss (Kaliss and Pressman, 1950). The quantity of evacetrapib in a volume of 8 µL of plasma was calculated using the average concentration in the plasma for each time point (in nmol/mL) multiplied by the volume. The literature shows that the conversion ratio of gram of brain to milliliter of brain is 1.04 mL for each gram(Leithner et al., 2010). Using the average concentration of evacetrapib (in nmol/g) in the brain and the average brain volume per gram, the average concentration (in nmol/mL) was calculated for each time point. From the calculated average concentration of evacetrapib in the brain (in nmol/mL), the quantity of evacetrapib (in nmol) in the brain for each time point was determined. Subtracting the amount of previously calculated average concentration of evacetrapib in the residual blood (in nmol/mL) to the average amount of evacetrapib in the brain for each time point (in nmol/mL), the average concentration of evacetrapib in the brain was corrected for residual blood (in nmol/mL and converted back to nmol/g).

Following the correction for the evacetrapib amount in the residual blood in the brain samples - using the area under the curve (AUC in nmol x h/mL) in the brain divided by the area under the curve in the plasma - the plasma-to-brain-tissue penetration ratio was calculated to determine the extent of which evacetrapib is detected in the brain depending on the initial concentration. The tissue penetration ratio allows to approximate how much evacetrapib is accumulated in the brain and whether it is consistent with dosage given to mice. The equation used to calculate this parameter is:

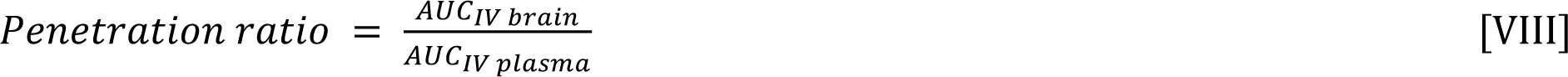

### Distribution to brain

To obtain more mechanistic information on the distribution of evacetrapib to the brain, a semi-physiological-based pharmacokinetic (PBPK) model was developed. First, a two-compartmental model (*i.e*., a plasma and a peripheral compartment) with a first order elimination rate was fitted to describe the compound’s kinetic in plasma (Mukherjee, 2022) (Figure 1). Second, a third compartment was added and connected to the plasma compartment (Figure 1). The parameters related to the brain compartment are presented Table 1. The brain compartment was subdivided into vascular and tissue sub-compartments, with mass balance differential equations describing a distribution in the tissue that can be limited by diffusion as follows (Shen, 2010):

**Figure 1:**
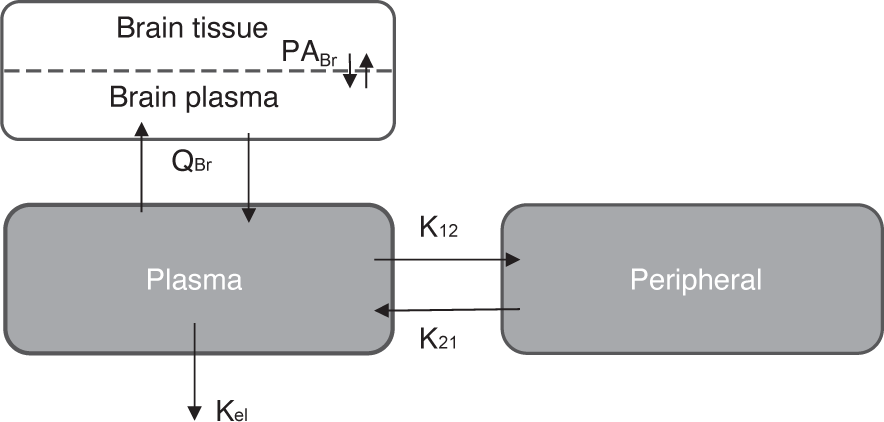
Schematic representation of the 3-compartment model simulation for the semi-PBPK model.

**Table 1:**
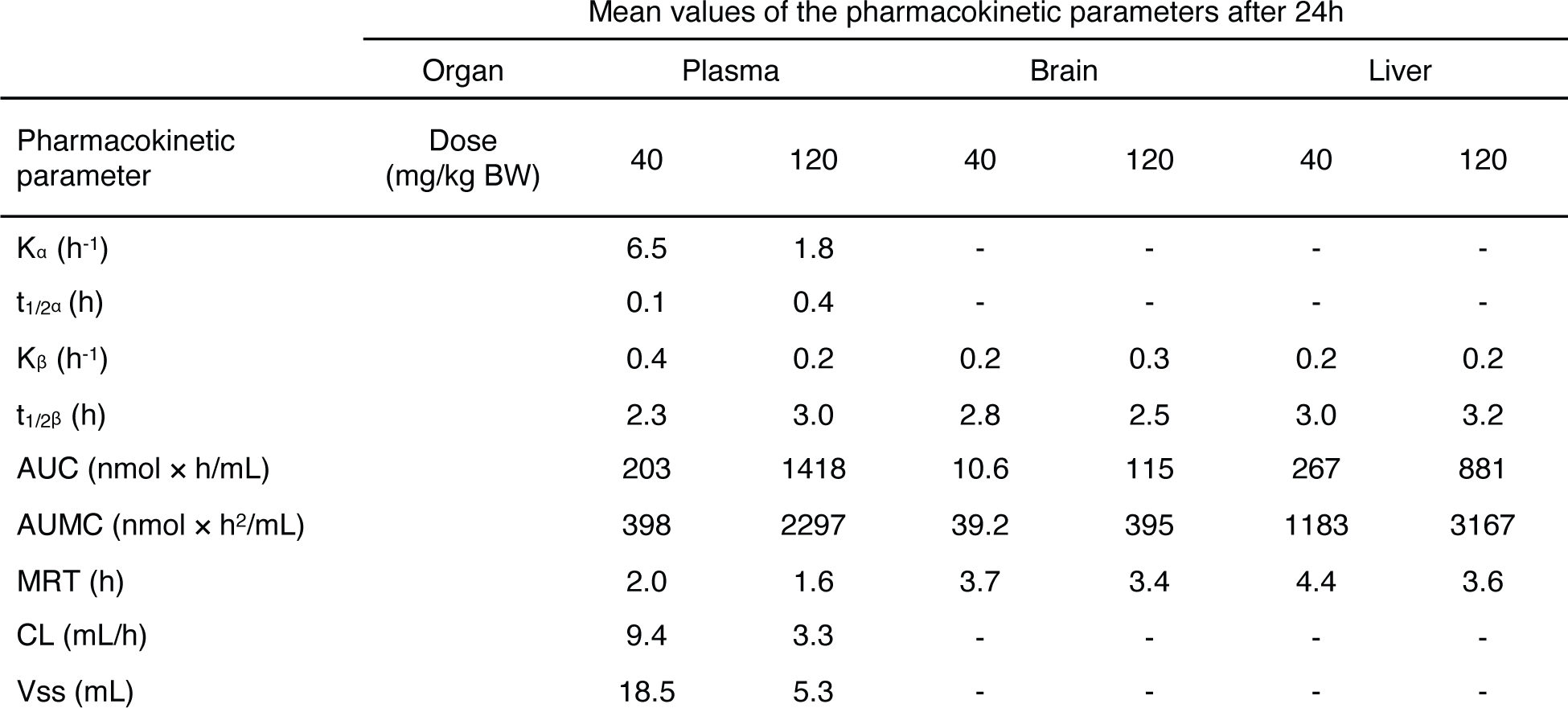
Pharmacokinetic parameters of evacetrapib in plasma, liver, and brain. *K: elimination rate constant; t_1/2_: half-life; AUC: area under the curve; AUMC: area under the first moment of the plasma concentration-time curve from time zero to infinity; MRT: mean residence time; CL: drug clearance; Vss: volume of distribution*.

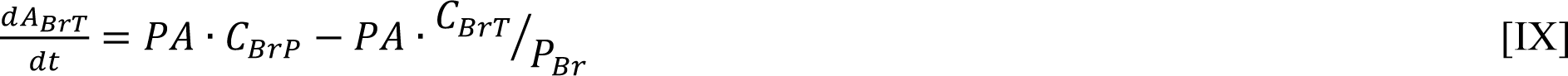

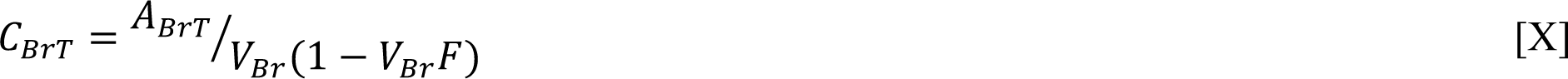

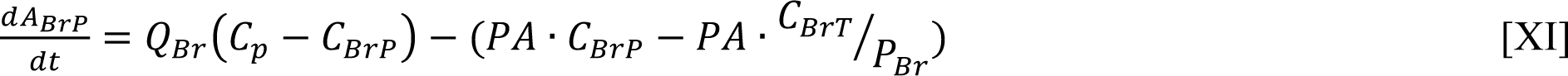

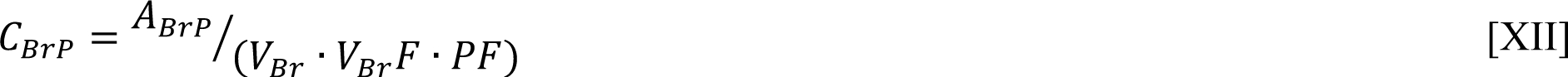

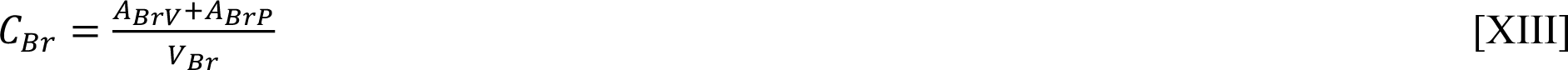

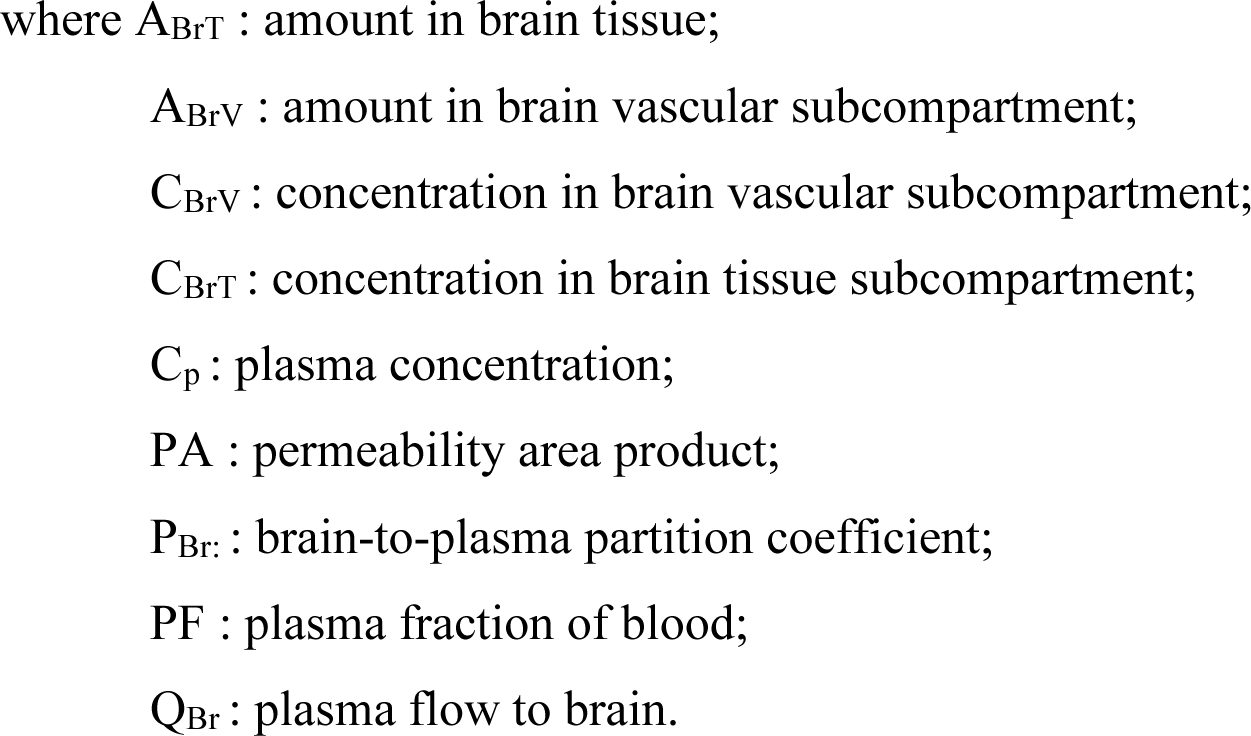

Using this semi-PBPK model and brain concentration data, several scenarios were tested to see what mechanisms best explain accumulation in the brain : 1) no permeability (i.e., PA=0 L/h); 2) a perfusion limited distribution (PA > 1,000,000 L/h) using the predicted P_Br_ (Poulin and Haddad, 2011); 3) a perfusion limited distribution with a low P_Br_ (fitted); 4) a diffusion limited distribution (fitted PA) with using the predicted P_Br_; 5) diffusion limited distribution (fitted PA) and a low P_Br_ (fitted).

## Results

Evacetrapib was shown to be tolerated up to 600 mg/kg BW/day in rats by oral gavage, with an oral absorption fraction of 25% (Simic et al., 2017; Takubo et al., 2014). We therefore chose a dose of 120 mg/kg BW as the highest intravenous dose and a 3-fold lower dose of 40 mg/kg BW for this pharmacokinetic study. Intravenous injections were chosen as the route of administration to bypass the barriers of absorption and to focus on distribution and elimination of the drug. Four male mice per time point were sacrificed and brain, blood and liver were collected according to the OECD guidelines 417 (OECD, 2010).

### Plasma elimination of evacetrapib

Peak plasma values were observed at the first sampling time point t=0 h followed by a rapid distribution phase to different tissues and then a second slower elimination phase from the plasma (Figure 2). Pharmacokinetic parameters were determined based on these blood time courses and are described in Table 1. The drug clearance (CL) from blood of 9.4 mL/h at the 40 mg/kg BW dose was much faster as compared to 3.3 mL/h at the 120 mg/kg BW dose. The mean residence time (MRT) of 2.0 h for the 40 mg/kg BW dose and 1.6 h for the 120 mg/kg BW dose remained similar. The steady state volume of distribution (Vss) of 18.5 mL was about three-fold greater at the 40 mg/kg BW dose as compared to 5.3 mL for the 120 mg/kg BW dose.

**Figure 2:**
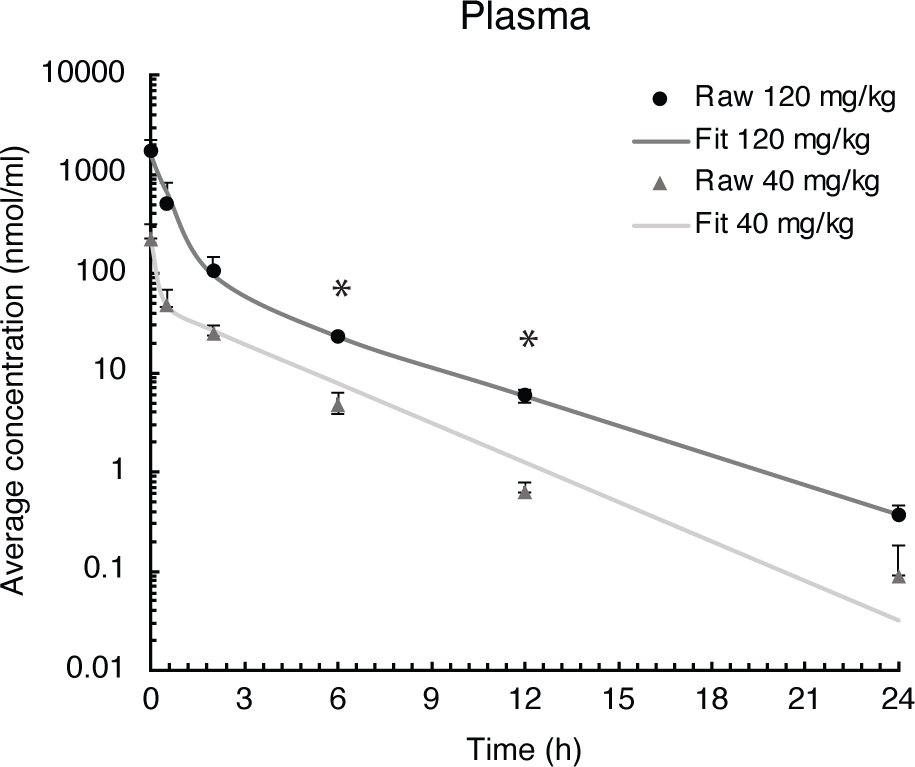
Evacetrapib concentration in plasma after a single intravenous injection of evacetrapib at 6 time points over a 24-h course. Symbols represent means ± s.e.m., n=4 per time point. The * depicts significant differences (p < 0.05) between the two doses.

### Elimination of evacetrapib in liver

The liver is the primary site for drug metabolism, responsible for the concentration, metabolism and excretion of the majority drugs that pass through the body. As such, evacetrapib quantification was determined in this tissue. For both doses in the liver, the time courses were very similar, showing a steady decrease of evacetrapib over time (Figure 3). Pharmacokinetic parameters were determined and described in Table 1. Results show similar elimination rates of 0.2 h^-1^ for the 40 mg/kg BW dose and 0.3 h^-1^ for the 120 mg/kg BW dose, with the slopes of both curves being parallel to each other. The elimination half-lives of 2.8 h and 2.5 h observed for the 40 mg/kg BW and 120 mg/kg BW, respectively, are of comparable value showing that both doses of evacetrapib are eliminated with the same speed. The MRT was similar for both concentrations (Table 1), though slightly higher for the 40 mg/kg BW indicating that the pharmacokinetic parameters in the liver remain unchanged both in the 40 mg/kg BW and 120 mg/kg BW conditions.

**Figure 3:**
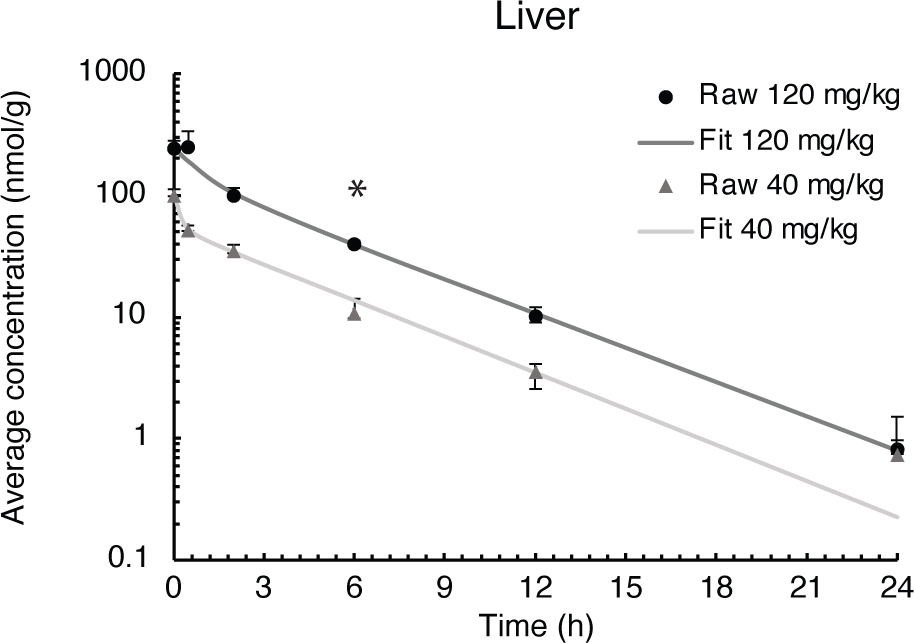
Evacetrapib concentration in liver after a single intravenous injection of evacetrapib. Symbols represent means ± s.e.m., n=4 per time point. The * depicts significant differences (p < 0.05) between the two doses.

### Evacetrapib pharmacokinetics in the brain

To assess whether evacetrapib can enter the brain, evacetrapib concentrations were quantified in this tissue. The time course for both doses shows that indeed evacetrapib enters the brain with a peak concentration after 0.5 h (Figure 4A). To have a more accurate reading of the concentration of evacetrapib in brain tissue, calculations were done to correct for the residual blood in the brain as shown in Table 2 and Figure 4B. For the 40 mg/kg BW dose at time 0 and 24 h, no evacetrapib was present in plasma and thus no correction was performed. At times 0.5, 2, 6, and 12 h, the average concentration of evacetrapib in the brain corrected for residual blood was determined as 1.5, 1.0, 0.5, and 0.1 nmol/g, respectively. For the 120 mg/kg BW dose, the average concentration of evacetrapib corrected for residual blood in the brain at time 0 h was 0 nmol/g. At times 0.5, 2, 6, 12, and 24 h, the average concentration of evacetrapib corrected for residual blood in the brain was determined as 16.7, 15.3, 5.0, 1.0, and 0.02 nmol/g, respectively, in a decreasing manner. This confirms the presence of evacetrapib in the brain tissue starting 30 minutes post-intravenous injection until 12 h post-injection. As reported in Table 1, the MRT values for both evacetrapib concentrations are similar with 3.7 h for the 40 mg/kg BW and 3.4 h for the 120 mg/kg BW dose showing that similar proportional amounts of evacetrapib enter the brain. Comparing the time courses of evacetrapib for not-corrected (Figure 4A) and corrected (Figure 4B) brain concentrations, the elimination rate constant k and half-lives remained similar, if not identical, for both doses. The MRT remained the same between the brain corrected and not corrected for residual blood in both concentrations as well.

**Figure 4:**
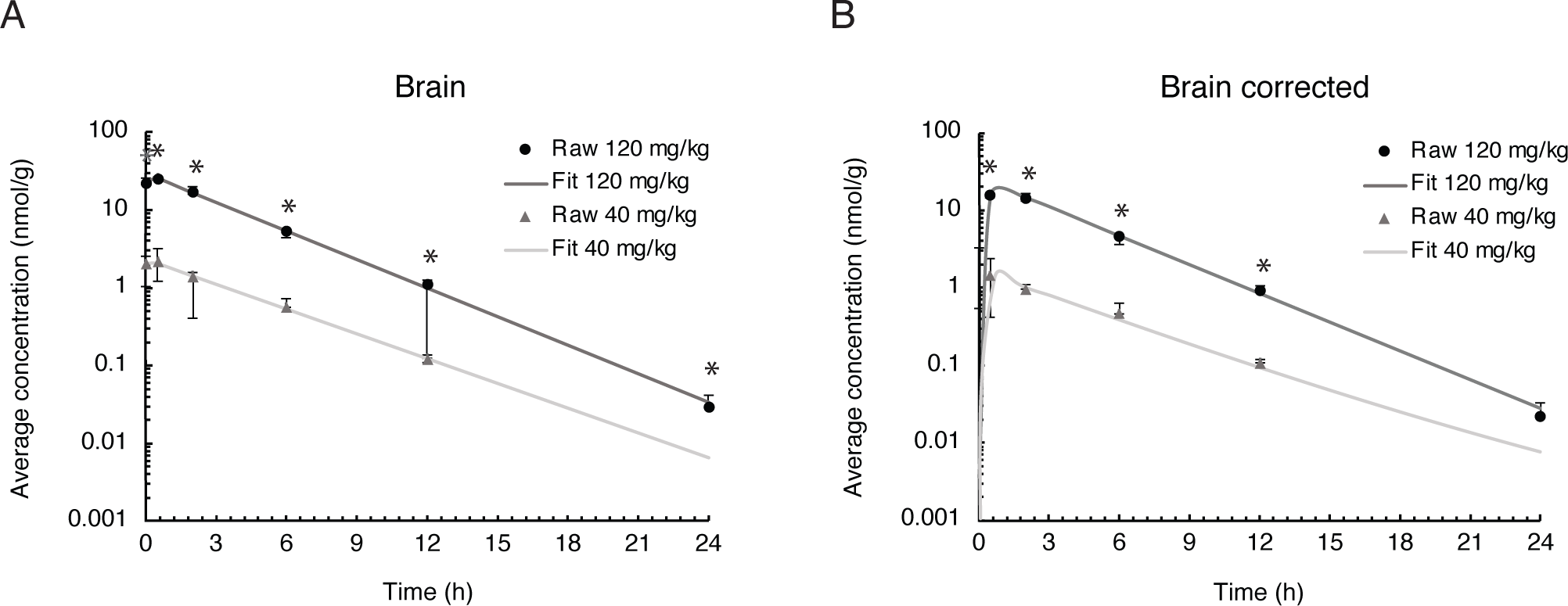
Evacetrapib concentration in the brain. A) Experimental values not corrected for residual blood. B) Experimental values corrected for residual blood. Symbols represent means ± s.e.m., n=4 per time point. The * depicts significant differences (p < 0.05) between the two doses.

**Table 2:** **Average evacetrapib concentration determined in the brain** over a 24-h *period after a 40 mg/kg BW and 120 mg/kg BW i.v. injection in hCETP tg mice comparing residual-blood correction to no correction*.

| | Concentration of evacetrapib in the brain for<br>40 mg/kg BW dose (mean ± SD) | | | Concentration of evacetrapib in the brain<br>for 120 mg/kg BW dose (mean ±SD) | | |
| --- | --- | --- | --- | --- | --- | --- |
|  | Measured | Calculated |  | Measured | Calculated |  |
| Time (h) | Average<br>concentration<br>(nmol/g) | Average<br>concentration<br>(nmol/g) | Standard<br>deviation<br>(nmol/g) | Average<br>concentration<br>(nmol/g) | Average<br>concentration<br>(nmol/g) | Standard<br>deviation<br>(nmol/g) |
| 0.00 | 2.05 | 0.00 | 0.54 | 22.2 | 0.00 | 3.30 |
| 0.50 | 2.21 | 1.53 | 1.00 | 25.8 | 16.7 | 0.97 |
| 2.00 | 1.41 | 1.03 | 0.15 | 17.1 | 15.3 | 2.60 |
| 6.00 | 0.57 | 0.50 | 0.17 | 5.36 | 4.96 | 0.21 |
| 12.00 | 0.12 | 0.11 | 0.01 | 1.11 | 1.00 | 0.14 |
| 24.00 | 0.00 | 0.00 | 0.00 | 0.03 | 0.02 | 0.01 |

### Tissue penetration ratio in the brain

The tissue penetration ratio calculated using the mean AUC (nmol × h/g) of the brain tissue divided by the mean AUC (nmol × h/g) of plasma and accounting for residual blood is described in Table 2 and yielded a ratio of 0.08 for the 40 mg/kg BW and 0.13 for the 120 mg/kg BW dose. Thus, the tissue penetration ratio shows a 1.63-fold increase from the low to the high evacetrapib concentration.

### Semi-PBPK model of evacetrapib entering the brain

We used a semi-physiologic three-compartment model to simulate the *in vivo* data (Figure 1). The parameters for the plasma and peripheral compartments were fitted to the plasma concentration at 40 mg/kg BW (Table 3 and Figure 5b) (Brown et al., 1997; Peyret et al., 2010). Then, four scenarios were simulated against the brain concentration data corrected for residual blood as explained in the method section. When assuming perfusion limited distribution, the model was not able to simulate the observed data (Figure 5a), even when we adjusted for the brain-to-plasma partition coefficient (P_Br_). However, modeling diffusion limitation in the brain resulted in a better fit of the simulation towards the observed brain concentrations when fitted for the permeability area product (PA) and P_Br_ (Figure 5a, dark grey curve compared to black circles).

**Figure 5:**
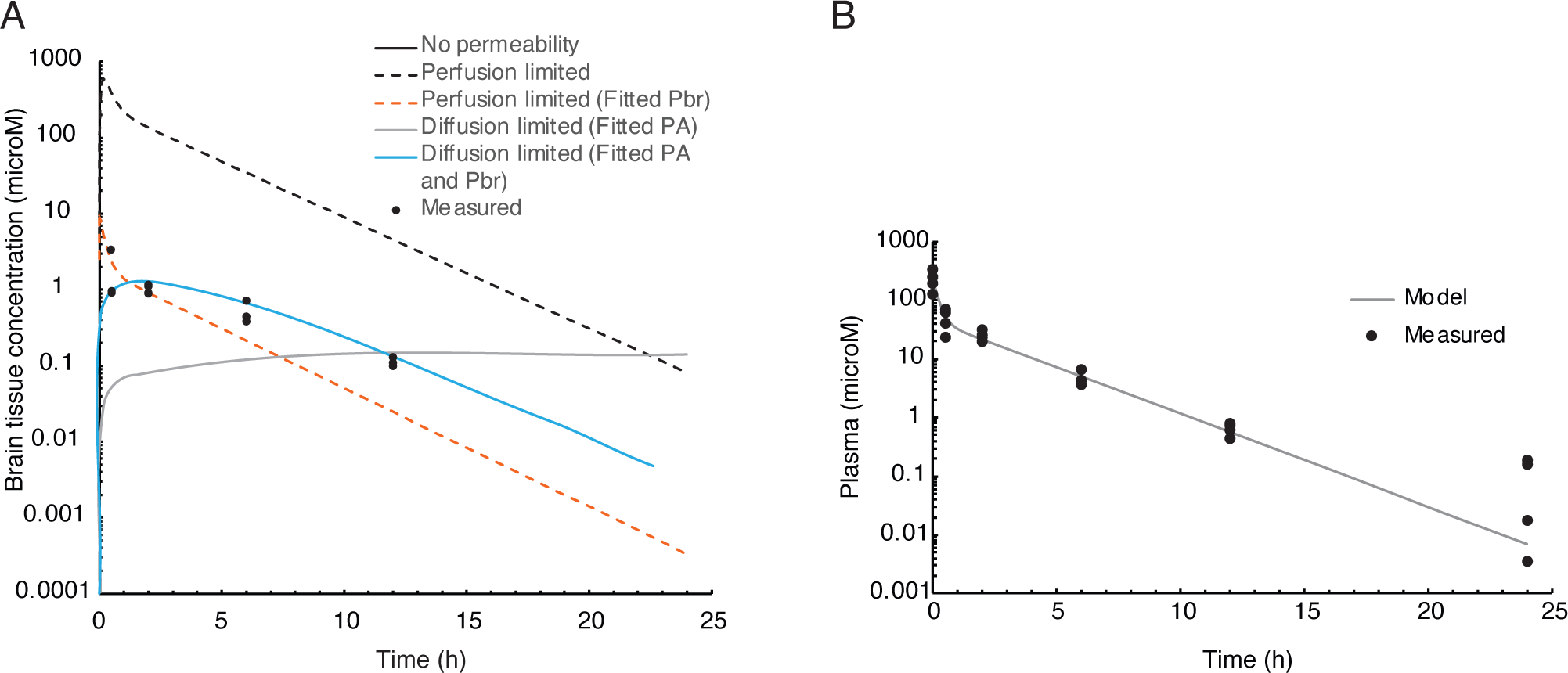
Semi-PBPK modeling of evacetrapib entering the brain. A) 3-compartment model simulation of the plasma concentration at 40 mg/kg. Experimentally determined values are in black circles. Different perfusion-limited models show different concentrations, with the best fit obtained with diffusion limitation and fitted PA and P_Br_. B) Parameters for the plasma and peripheral compartments fitted to the plasma concentration at 40 mg/kg BW.

**Table 3:** Parameter values for the semi physiologically-based pharmacokinetic model to simulate accumulation in the brain.

| Two-compartment model | Abbreviation | Value | Source |
| --- | --- | --- | --- |
| Apparent plasma volume (L) | $V_{\text{plasma}}$ | 0.01 | Fitted |
| Plasma to peripheral transfer constant ($\text{h}^{-1}$) | $K_{12}$ | 2.4 | Fitted |
| Peripheral to plasma transfer constant ($\text{h}^{-1}$) | $K_{21}$ | 1.21 | Fitted |
| Elimination constant ($\text{h}^{-1}$) | $K_{\text{el}}$ | 1.39 | Fitted |
| **Added brain compartment** |  |  |  |
| Brain volume fraction (fraction of body weight) | $V_{\text{BrF}}$ | 0.0165 | Brown et al. <sup>30</sup> |
| Vascular volume fraction (fraction of compartment) | $V_{\text{Brvf}}$ | 0.03 | Brown et al. <sup>30</sup> |
| Cardiac output (L/h) | $Q_{\text{C}}$ | 0.8388 | Brown et al. <sup>30</sup> |
| Plasma flow in brain as fraction of cardiac output | $Q_{\text{BrF}}$ | 0.033 | Brown et al. <sup>30</sup> |
| Brain:plasma Partition coefficient | $P_{\text{Br}}$ | 6.9 | Estimated <sup>31</sup> |
|  |  | 0.06 | Fitted |
| Permeability area product (L/h) | $PA$ | $1.1 \times 10^{-5}$ | Fitted |

## Discussion

We investigated the distribution of the CETP inhibitor evacetrapib in hCETPtg mice, and the major finding of this study is that evacetrapib does enter mouse brain tissue. We detected evacetrapib in the brain 0.5 h after intravenous injection at already 40 mg/kg BW in a nonlinear function. Evacetrapib inhibits CETP in human plasma with an IC_50_ of 36 nM (Cao et al., 2011). We detected evacetrapib in the brain at peak (after correction for residual blood) of 1.5 µM at 40 mg/kg BW, and of 16.7 µM at 120 mg/kg BW dose, thus approximately 40-fold or 450-fold, respectively, above the plasma IC_50_, indicating that evacetrapib administration could indeed inhibit cerebral CETP, potentially even at lower doses. Our semi-PBPK compartmental model shows that evacetrapib does not rapidly diffuse across the blood brain barrier indicating that the mechanism of evacetrapib distribution to the brain may rely on facilitated transport mechanisms.

Dysregulation of the cholesterol metabolism has been associated with various diseases such as atherosclerosis*^7^* and Alzheimer’s disease (Liu et al., 2020; Vance, 2012). An important protein regulating the distribution of cholesteryl esters in the blood is CETP, which has been linked to Alzheimer’s disease through polymorphisms (Lythgoe et al., 2015; Rodriguez et al., 2006; Sundermann et al., 2016). Though the function of CETP has mainly been established in the periphery, cholesteryl transfer activity has also been reported in the central nervous system (CNS) and in the cerebrospinal fluid (CSF) (Albers et al., 1992; Yamada et al., 1995). While it has been suggested that plasma cholesterol does not have a direct effect on brain cholesterol levels (Bjorkhem and Meaney, 2004), it remains unclear whether it is CETP activity in the CNS or in the blood that plays a role in dementia. Lipoprotein particles of the brain differ from those of the periphery as LDL-like particles do not exist in the brain, but rather small HDL-like particles which are only incompletely characterized (Vitali et al., 2014). These HDL-like particles transport cholesterol from astrocytes to neurons (Borras et al., 2022; Ioannou et al., 2019; Turri et al., 2022). Since CETP remodels HDL particles (Nicholls et al., 2015), a potential benefit of CETP inhibition could stem from promoted cholesterol delivery to neurons.

In a given model organism, it is therefore necessary to assess CETP activity and LDL levels *in vivo*. However, unlike humans, mice do not express endogenous CETP, making hCETPtg mice a more representative animal model in terms of peripheral lipoprotein dynamics. Indeed, while wild type mice have very low levels of endogenous LDL, hCETP tg mice have an LDL profile that is much more comparable to humans due to CETP activity (Steenbergen et al., 2010). We previously found that hCETPtg mice have approximately 22% higher cholesterol content in their brains compared to wild type mouse brains (Oestereich et al., 2022). The elevated cholesterol content in the brain and in the periphery of hCETPtg mice could differentially affect the distribution and elimination kinetics of the very lipophilic CETP inhibitors (evacetrapib has an octanol to water partition coefficient of logP 7.56) (Small et al., 2015). Due to these fundamental physiological differences, pharmacokinetic parameters of a CETP inhibitor should be assessed in hCETPtg mice rather than wild type mice. Thus, hCETPtg mice present a good model for testing CETP inhibitors in the context of drug repurposing for Alzheimer’s disease. Remarkably, several proteins associated with Alzheimer’s disease are linked to cholesterol. For example, the central protein in Alzheimer’s disease, the amyloid precursor-protein (APP), contains cholesterol-binding motifs in its transmembrane sequence (Barrett et al., 2012). Further proteins encoded by gene variants identified by genome-wide associated studies carry functions related to cholesterol including the most prominent APOE, CLU, SORL1, PICALM, BIN1, SORT1, and ABCA1 (Bellenguez et al., 2022). As such, CETP may have an indirect modifying role in the Alzheimer’s disease by affecting the cholesterol of the brain. Collectively, further investigations are necessary to address the potential impact of CETP inhibition in ameliorating Alzheimer’s disease symptoms by reducing cerebral cholesterol.

Only one other CETP inhibitor, dalcetrapib developed by Hoffman-La Roche, was shown to have the capacity to cross the blood-brain barrier of rats (Takubo et al., 2014). In comparison to our study, rats received 10 mg/kg BW dalcetrapib intravenously, which reached a 2-fold lower concentration in the brain after 0.5 h than 40 mg/kg BW evacetrapib in mice. Conversely, at 24 h post-injection, the dalcetrapib concentration was >20-fold higher than the evacetrapib concentration in mice injected with 40 mg/kg BW indicating that, though evacetrapib may enter the brain tissue faster than dalcetrapib, it has a lower capacity to accumulate and is eliminated faster than dalcetrapib. It should be noted that anacetrapib, another CETP inhibitor developed by Merck, has been shown to accumulate in adipose tissues (Krishna et al., 2017). In contrast, continuous administration of evacetrapib does not accumulate in adipose tissues (Nurmohamed et al., 2022; Small et al., 2015), avoiding unwanted storage of the drug in the body. To our knowledge, whether the CETP inhibitor evacetrapib accumulates in the brain with long-term use has not yet been evaluated. It should also be noted that the doses administered for this pharmacokinetic study in mice are higher than those tested in clinical trials in humans (Nair and Jacob, 2016; Nicholls et al., 2011), while lower doses will likely still reach the brain at biologically relevant concentrations.

One last factor we considered in this study was the mechanism by which evacetrapib is transported into brain tissue. To that effect, toxicokinetic parameters were derived from the blood and brain time courses, which indicate that the drug possibly requires some sort of transporter to reach peripheral compartments, such as the brain, and in order to clear from brain tissue. The difference in tissue penetration ratio as a function of the dose indicates that a higher concentration of evacetrapib has greater capacity to be detected in brain tissue. A possible explanation could be that evacetrapib enters the brain faster than it is cleared out of it, leading to a buildup in brain tissue. The compartmental model developed (Shen, 2010) suggests that diffusion-limited uptake occurs in the brain and that the P_Br_ was very low, which may indicate that evacetrapib diffuses into the brain inefficiently and would require the use of transporters. To determine which mechanisms are involved in evacetrapib distribution to the brain, a semi-PBPK model was developed (parameter values are in Table 3; Figure 5). To this end, we first aimed to determine the brain:plasma partition coefficient of evacetrapib using rapid equilibrium dialysis. Our preliminary data showed that the molecule did not appreciably diffuse across the porous membrane; only 2% of evacetrapib diffused after 24 h in the system consisting of plasma on both sides of the membrane. On the one hand, this may be in line with our *in vivo* observations considering that only a small fraction of evacetrapib crossed the blood-brain-barrier. However, the slow diffusion in our *in vitro* model is incongruent to the concentrations of evacetrapib found in the brain 2.2 h following intravenous injections. Thus, the implication of transporters cannot be excluded.

Epidemiological studies indicated that CETP plays a disease-modifying role in Alzheimer’s disease (Barzilai et al., 2003; Rodriguez et al., 2006; Sun et al., 2013). While the exact mechanism remains unclear, CETP activity may promote Alzheimer’s disease through modifying cholesterol content, distribution, storage, or metabolism, which could potentially be prevented by CETP inhibitors such as evacetrapib. To set the grounds to assess CETP inhibitors for potential drug repurposing in Alzheimer’s disease, we showed here that evacetrapib reaches brain tissue fast after intravenous injection in hCETPtg mice. Furthermore, we demonstrate that evacetrapib crosses to the brain, possibly through the actions of a transporter or other facilitating mechanism. Thus, CETP activity in the brain could be pharmacologically reversed, which may carry the potential to delay or ameliorate Alzheimer’s disease.

## Acknowledgements

We thank the CMARC Animal Health Technicians for the drug treatments in mice.

## Author contributions

JP wrote the manuscript draft, made the figures, bread mice and collected samples after injections. JC modeled the pharmacokinetic parameters. JP and JC performed the pharmacokinetic analysis. DD conducted and analysed evacetrapib detection by mass spectrometry. SJR, SE and FO bread mice and collected mouse samples after injections. SH designed and completed the semi-PBPK model. SH, MB, and LMM designed, coordinated, and supervised the study. All authors revised the manuscript.

